# The compound topology of a continent-wide interaction network explained by an integrative hypothesis of specialization

**DOI:** 10.1101/236687

**Authors:** Gabriel Moreira Felix, Rafael Barros Pereira Pinheiro, Robert Poulin, Boris R. Krasnov, Marco Aurelio Ribeiro de Mello

## Abstract

Is there a prevalent pattern among interaction networks: nestedness or modularity? Must consumers always trade-off generalism for average performance in resource exploitation? These two questions have been addressed in various systems, with contradictory results. A recent integrative hypothesis combines both questions within a common theoretical framework, proposing that ecological specialization is structured by different prevailing processes in smaller and larger network units. This should produce both a compound interaction network, formed by internally nested modules, and a scale-dependence on the relationship between consumer performance and generalism. Here, we confirm both predictions in a large dataset on host-parasite interactions. We show that modules indeed constrain nestedness at the whole network level, and that the relationship between parasite generalism and performance on their hosts changed from negative at large to positive at small scales. Our results shed light on both debates, and provide some clues to their integration and solution.

## INTRODUCTION

Darwin’s “tangled bank” of species interactions is one of the most complex phenomena in nature. In the past decades, ecologists have imported analytical tools from network science (Barabási 2016) to disentangle this complexity (Bascompte & Jordano 2013). Despite the progress made in describing pervasive patterns and underlying mechanisms (Vazquez *et al*. 2009), some aspects of the architecture of ecological networks remain controversial, begging for further investigation (Dormann *et al*. 2017). One of those unanswered questions concerns what should be the predominant topology among ecological networks: nestedness or modularity (Thebault & Fontaine 2010).

By adapting the biogeographic concept of nestedness (Atmar & Patterson 1993) to interaction matrices, Bascompte *et al*. (2003) showed that several plant-animal networks have a nested topology, with the interactions of specialists tending to be subsets of the interactions of generalists. Later studies also found nestedness in many other mutualistic (Ollerton *et al*. 2003, 2007; Guimaraes *et al*. 2006) and antagonistic networks (Vázquez *et al*. 2007; Graham *et al*. 2009). Meanwhile, another topology, modularity, has also been widely reported (Olesen *et al*. 2007; Dupont & Olesen 2009; Mello *et al*. 2011; Krasnov *et al*. 2012). Since a modular network is one composed of modules of species that interact more frequently with one another than with other species of the same network, forbidden interactions between modules should constrain nestedness. Therefore, the two topologies seem to be mutually exclusive. Nevertheless, some ecological networks present high scores of both nestedness and modularity, and a positive relationship between these two topologies has even been reported (Fortuna *et al*. 2010).

Thus, a new question arose: how can a network be both modular and nested at the same time? A possible answer states that a dual nested-modular structure would arise if each topology predominates at different structural scales of the network (Lewinsohn *et al*. 2006). Specifically, the authors suggested that some plant-animal networks are modular at the scale of the entire network, but their modules are internally nested. This kind of multi-scale architecture was named a compound (or combined) topology. Later, some empirical studies found evidence of compound topologies in pollination (Bezerra *et al*. 2009), seed dispersal (Sarmento *et al*. 2014), and phage-bacteria networks (Flores *et al*. 2013). In addition, theoretical studies confirmed this topology in simulated host-parasite networks (Beckett & Williams 2013; Leung & Weitz 2016).

Recently, an “integrative hypothesis of specialization” (IHS) was advanced, which proposed a mechanism by which a compound topology might emerge in ecological networks (Pinheiro *et al*. 2016). If its logic is correct, this hypothesis would also help to solve another important ecological controversy: what is the expected relationship between the resource range (generalism) of a given consumer and its average performance in exploring these resources (Futuyma & Moreno 1988)? The IHS states that this relationship should change with scale, from negative across the network as a whole to positive within each module, which should lead to the emergence of an interaction network formed by internally nested modules. The rationale of the IHS is briefly described in Box 1 and Fig. 1.

**Figure 1:**
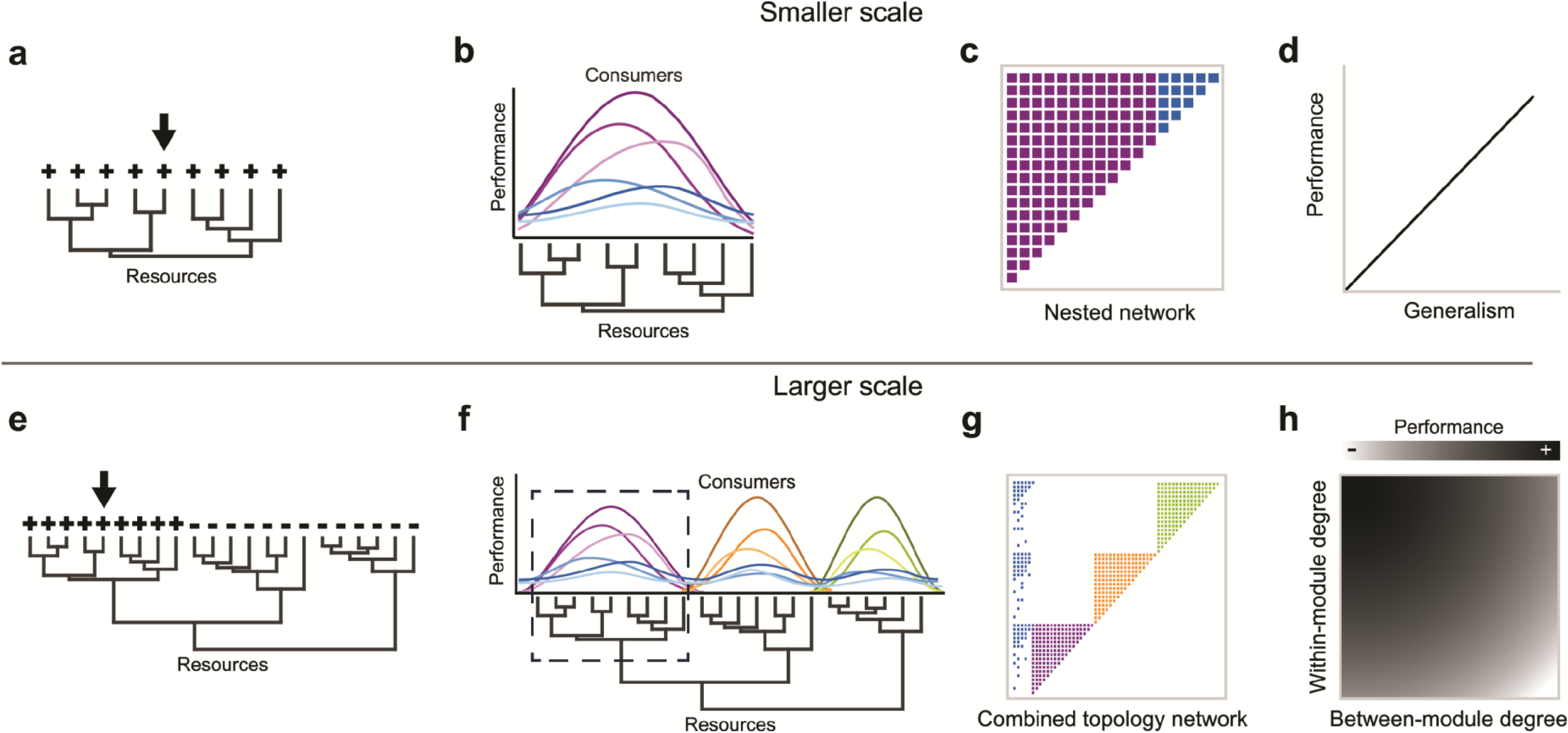
The integrative hypothesis of specialization. Explanations are given in Box1.

Despite their potential to help solve two important ecological debates, and to improve our understanding of the structure of ecological communities, neither the compound topology nor the IHS has been widely tested. Here, we address these two issues in an extensive host-parasite data set composed of flea-mammal interactions in 15 Palearctic regions. Since this data set was collected at large phylogenetic and geographic scales, it is a good model to test the relationship between generalism and performance, and also the existence of a compound topology.

First, we adapted the method described by Flores *et al*. (2013) and developed a general framework to test nestedness at different network scales. Then, we tested whether the flea-mammal networks have a compound topology. This question was addressed in both the global network (formed by pooling together interactions reported in different regions) and the local networks (formed by interactions reported for each region). Second, we tested whether the relationship between host range (generalism) and performance in fleas is scale-dependent, changing from positive within clusters of similar resources (within each module) to negative between clusters (between modules). This second question was addressed using only the local networks, since constraints in the global network are mainly geographic (two species need to co-occur in the same site to interact with one another) and would not reflect trade-offs in specialization. Our results shed light on both debates, and provide some clues to their integration and solution.

## METHODS

### Data set

We used an extensive host-parasite data set that has been analyzed in several studies on ecological interactions (*e.g*, Krasnov *et al*. 2004, 2008; Vázquez *et al*. 2007; Fortuna *et al*. 2010). It is composed of dozens of flea-mammal interaction matrices sampled all around the world, from which we selected 15 Palearctic regions (see Table S1 in Appendix S2) to maximize two parameters: the size of the matrix (at least 10 parasite species and 10 host species per region), and the number of hosts sampled (more than 1,000 individual mammals per region).

The global matrix with all 15 regions pooled has a size of 263 species (nodes: 161 fleas and 102 mammals), and contains 1,200 interaction records (links). The local networks have an average size of 45.06 ± 12.64 species (mean ± standard deviation), with 26.26 ± 9.42 fleas and 18.8 ± 4.79 mammals, and contain on average 129.6 ± 57.22 interaction records (Appendix S2: Table S1).

Furthermore, the global matrix and some local matrices produced networks with more than one component, *i.e*., a cluster of species totally separated from the other nodes of the network. In most of these networks, there is a giant component comprising most of the network nodes and one or few minor components, each one including a small number of nodes. The analyses below, at global and local scales, were carried out by using only the respective binary version of the largest component of each matrix.

## Network topology

### Modularity

The first step to test for a compound topology is to unfold the modular structure of the network. We did this computing the Barber modularity (Q) (Barber 2007) optimized by the DIRTLPAwb+ algorithm (Beckett 2016), through the *computeModules* function of the *bipartite* package (Dormann *et al*. 2008) for R (R Development Core Team 2017). Modularity (Q) varies from 0 to 1, and the algorithm reveals also the number and composition of the modules found in the network.

### Nestedness

A nested matrix has its interactions arranged in a particular way: the interactions of the least connected species are proper subsets of the interactions of the more connected species (Ulrich *et al*. 2009). NODF is a metric that aims to synthesize this pattern in a single number (Almeida-Neto *et al*. 2008). In its default procedure, a NODF score is computed for each pair of species (independently for consumers and resources, i.e., rows and columns of the interaction matrix) and, then, averaged to calculate the NODF score of the whole matrix. This procedure implicitly assumes that nestedness is evenly distributed in the matrix. However, as pointed out by the authors of NODF themselves, it is important to “*explore whether nestedness is a general pattern of the community or derives from some particular species subsets*” (Almeida-Neto *et al*. 2008). If different species subsets of the matrix have different degrees of nestedness, an overall NODF is not an appropriate summary of the matrix structure (Gotelli & Ulrich 2012). It turns out that this is exactly the case if the network has a compound topology, where nestedness between pairs of species of the same module should be much higher than nestedness between pairs of species of different modules.

In order to solve this problem, we adapted the method described in Flores *et al*. (2013), and averaged nestedness independently between pairs of species of the same module (NODF_SM_), and between pairs of species of different modules (NODF_DM_), and compared those values with those expected by species degrees under two scenarios (null models): in the absence of a modular structure (free null model) and in the presence of a modular structure (restricted null model). The rationale behind using these two null models is explained below, while the detailed instructions for performing both null models are presented in Appendix S1.

### Predictions

If nestedness and modularity coexist at large network scales as two sides of the same coin, we expected nestedness between species of different modules (NODF_DM_) to be equal to or higher than expected by their degrees (*i.e*., the free null model). Otherwise, if modularity constrains nestedness between pairs of species of different modules, we expect NODF_DM_ to be smaller than expected by their degrees.

Notice, however, that NODF between pairs of species of the same module (NODF_SM_) will be higher than expected by species degrees whether or not modules are internally nested. This would happen since, by definition, species of the same module share more interactions with one another than expected by their degrees, regardless of those interactions being nested or not. Hence, a NODF_SM_ value higher than expected by the free null model is a necessary, but not sufficient condition, for a network to have a compound topology. This is the reason why we need the restricted null model – which also conserves the modular structure of the original matrix when generating the null matrices (Additional Information 1) –, to test whether NODF_SM_ is higher than expected *given* the modular structure. On the one hand, if the network is formed by modules that are not internally nested, NODF_SM_ should be higher than expected by the free null model, but lower than expected by the restricted null model. On the other hand, if the network is formed by internally nested modules (*i.e*., a compound topology), NODF_SM_ should be higher than expected by both the free and restricted null models.

But why did we not individualize each module and, then, test its nestedness independently? Although this would be a valid procedure to test NODF_SM_, it would not allow to test whether interactions between pairs of species of different modules (NODF_DM_) are more nested than expected *given* the constraints imposed by the modules. This can be done by comparing the observed NODF_DM_ with that expected by the restricted null model.

### Z-Score

For each main component of the 16 networks (the global network and 15 local networks), we generated 1,000 random matrices using the free null model and another 1,000 matrices using the restricted null model. Next, for each random matrix, we computed its overall NODF and decomposed it into NODF_SM_ and NODF_DM_ using the observed partitions of their corresponding real network.

Finally, for all combination of matrices (16 in total: 1 global and 15 local), null models (2: free and restricted), and NODF metrics (3: NODF, NODF_SM_ and NODF_DM_), a Z-score was calculated as Z = [Value_obs_ – mean(Value_sim_)] / σ(Value_sim_), where Value_obs_ is the observed value of the metric and Value_sim_ represents the values of the metric in the randomized matrices. Observed and expected modularity values were also compared using Z-scores, but only for the free null model, as it does not make sense to compare observed and expected modularities with a null model that fixes the modules.

Nestedness and modularity standardized by null models will be called relative nestedness and relative modularity, respectively. For simplicity, they will be represented here as Z_F_ or Z_R_, depending on the null model, followed by the metric name (e.g, Z_F_Q and Z_F_NODF_SM_ represent, respectively, relative modularity and relative nestedness between pairs of species of the same module, when standardized by the free null model).

Our goal was to see how modularity and nestedness interact with each other in a continuous way. Therefore, in all analyses we used the original Z-scores, without classifying them as significant and non-significant.

### Matrix plotting

The interaction matrices were reorganized to maximize between- and within-module nestedness as done in previous studies (Flores *et al*. 2013, 2016). Briefly, we first reordered the matrix rows and columns by degree without disrupting its modular structure and, then, permuted the modules in order to find the arrangement of modules which maximizes the overall NODF of the matrix. This procedure facilitates the visualization of a compound topology, if one exists.

## Specialization versus performance at different scales

### Performance index

In the host-parasite literature, the performance of a parasite in a host is usually quantified indirectly through some metric assumed to reflect it: *e.g*, prevalence, intensity, or abundance (Poulin 2007). We chose abundance: the average number of individual fleas per individual mammal (calculated including infected and uninfected hosts). This choice is justified as abundance is considered a good measure of performance in host-parasite systems (Krasnov *et al*. 2006), since it integrates intensity of infestation and prevalence in a single metric (abundance = intensity of infestation *times* prevalence), measuring different aspects of parasite performance. While intensity of infestation is the average number of individual parasites per infected individual host, prevalence is the proportion of infected individuals in the host population.

### Generalism within modules and between modules

For each flea species, specialization within and between modules was measured through its cartographic position in the network: its within-module degree (Z, which should not be confused with the Z-score of the null models) and participation coefficient (P) (Guimerà & Amaral 2005).

These two metrics define the functional role of a species in a network, and they are respectively related to the number of interactions a species makes with other species of its own module and with species of other modules. Z and P values were calculated independently for each local network.

### Mixed models

We used mixed models (Bolker *et al*. 2009) to test whether flea performances are positively correlated with their within-module degrees (Z) and negatively correlated with their participation coefficients (P). Linear mixed models (LMMs) were built by the *lmer* function of the *lme4* package (Bates *et al*. 2015), and fitted by restricted maximum likelihood (REML).

We used the log-transformed abundance of each parasite species in each host species in each region as the response variable, Z and P values for each flea species in each region as the explanatory variables, and host species, parasite species and region as crossed random factors. We decided to use parasite abundances per host species, rather than average it between all hosts exploited by a flea, to control for host characteristics known to affect abundance (e.g., carrying capacity, susceptibility, and richness of parasite fauna) (Krasnov *et al*. 2005). Averaging would also decrease the power of the analysis (Hopkins 1982; Schank & Koehnle 2009).

In addition, as pointed out in Box1 (see also Fig. 4 in Pinheiro *et al*. (2016)), in local networks composed of very similar resources, we should not expect to find either a negative relationship between abundance and P or modules. In those networks, the modules recovered by the DIRTLPAwb+ algorithm will be spurious, not imposing constraints to interactions, and the measured NODF_DM_ should be higher than expected by species degrees (the free null model). To test this prediction, we included an interaction between Z_F_NODF_DM_ and the fixed factors of the model (P and Z). We expect Z_F_NODF_DM_ not to have an influence on the effect of Z on abundance, but to influence the effect of P. Specifically, we expected that the effect of P on abundance should be negative only in local networks in which the modular structure constrains nestedness between pairs of species of different modules, that is, in local networks with negative values of Z_F_NODF_DM_.

We used backward stepwise regression to select fixed and random effects, following the procedure suggested by Bolker *et al*. (2009). We used the *anova* function of the *stats* package to perform a likelihood ratio (LR) test on the random effects (to which the models are refitted with maximum likelihood) and, then, used the *Anova* function of the *car* package to perform Wald *X*^2^ tests on the fixed effects. To tell apart the variance explained by either the fixed or random factors in the minimal selected model, we used the *r.squaredGLMM* function of the *MuMIn* package for R to compute both marginal and conditional R squared (Nakagawa & Schielzeth 2013). The confidence intervals of the parameters were obtained by bootstraping using the *confint.merMod* function of *lme4* package. The confidence interval of the conditional effect of P given Z_F_NODF_DM_ was also computed by simulation using the *interplot* function of *interplot* package.

## RESULTS

### Topology

#### Global network

The global network presented higher modularity (Z_F_Q = 51.13) and overall nestedness equal (Z_F_NODF = 0.39) to that expected by the free null model. However, the observed scores of nestedness were much higher between pairs of species of the same module than between pairs of species of different modules (NODF_SM_ = 45.47, NODF_DM_ = 9.85). In addition, as expected if the modules constrain nestedness between species of different modules, NODF_DM_ was smaller than expected by the free null model (Z_F_NODF_DM_= −13.46) (Fig. 2a), but equal to expected by the restricted null model (Z_R_NODF_DM_ = 0.22) (Fig. 2b). Finally, nestedness between pairs of species at the same module was higher than expected by both null models (Z_F_NODF_SM_ = 53.89, Z_R_NODF_SM_ = 22.08) (Fig. 2a–b).

**Figure 2:**
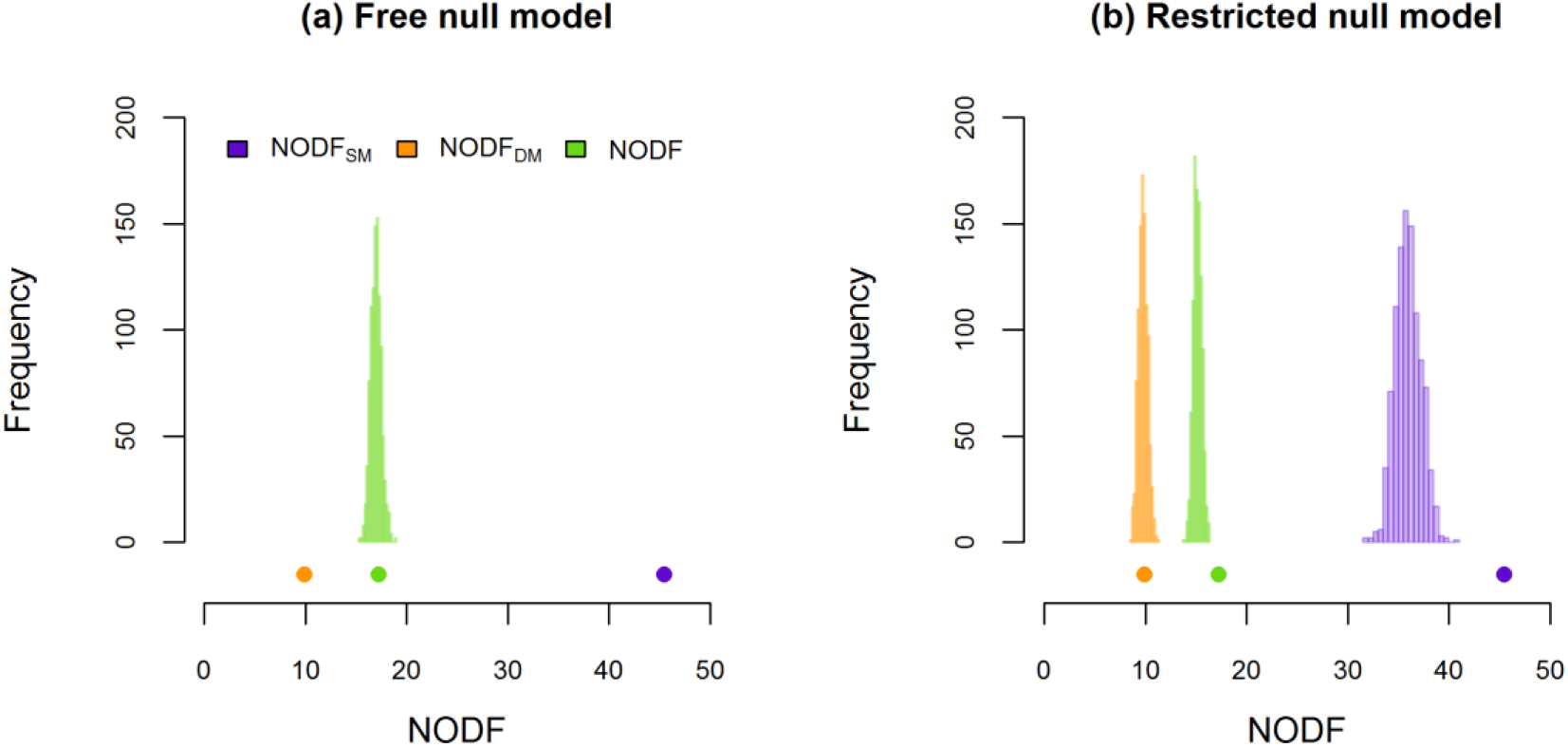
Observed values (dots) of NODF, NODF_SM_, and NODF_DM_ in the global network contrasted with the values expected by species degrees (distributions) in the absence (a: free null model) or presence (b: restricted null model) of a modular structure. NODF: overall nestedness. NODF_SM_: nestedness between pairs of species of the same module. NODF_DM_: nestedness between pairs of species of different modules. As expected if the global network has a compound topology, NODF_SM_ is higher than expected by both null models, and NODF_DM_ is smaller than expected by the free null model and equal to that expected by the restricted null model.

Those results strongly support the hypothesis that the global flea-mammal network has a compound topology, which can be easily seen when we plot the interaction matrix maximizing nestedness without disrupting its modular structure (Fig 3).

**Fig. 3:**
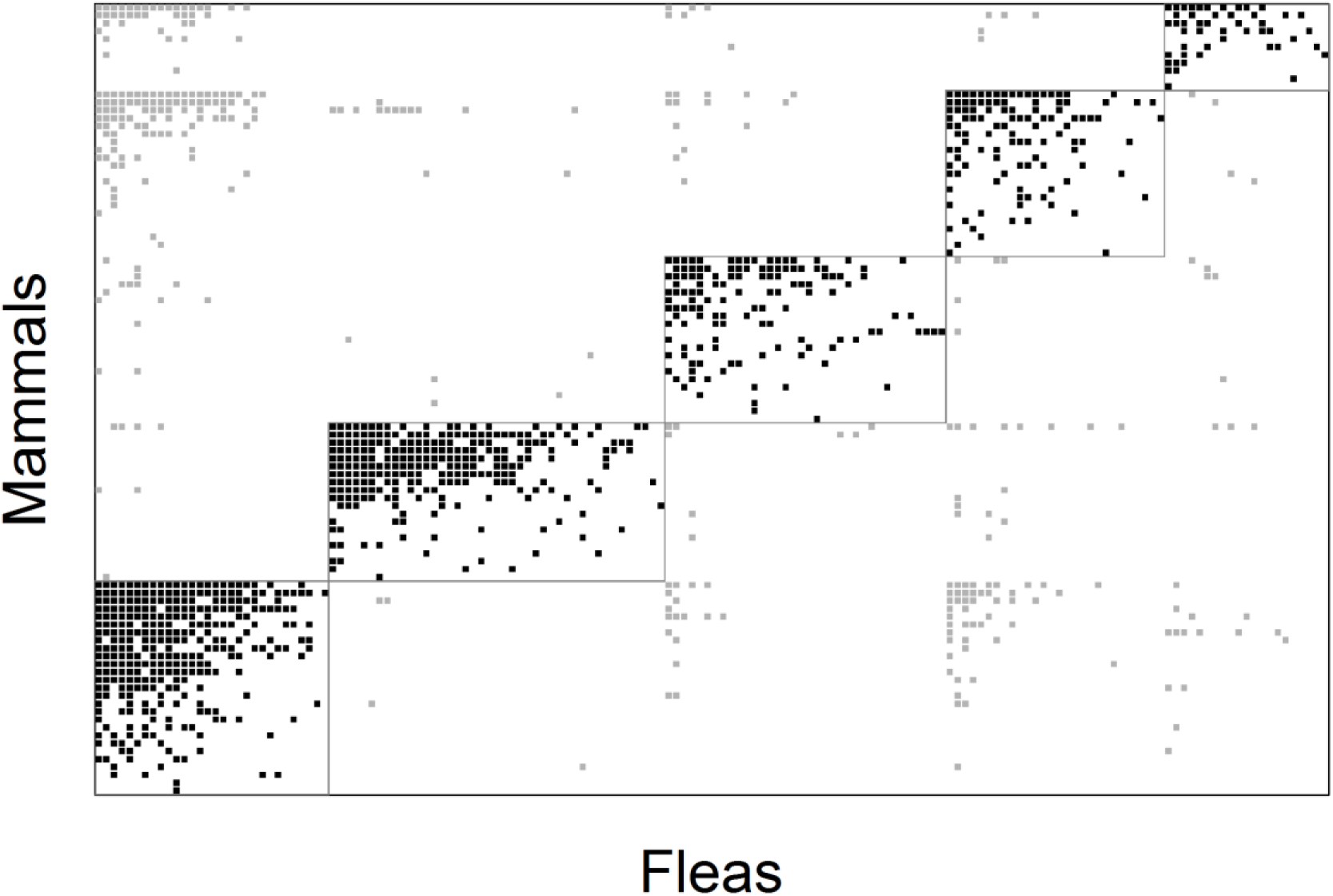
Interaction matrix reorganized to maximize between- and within-module nestedness without disrupting the modular structure of the network. Interactions within modules (delimited by boxes) are showed in black, while those outside modules are showed in gray. Flea species are represented in rows and mammals in columns. The compound topology of the global network is evident.

#### Local networks

Nestedness and modularity varied widely among local networks, which, in general, were more nested (Z_F_NODF = 2.03 ± 1.94) than modular (Z_F_Q = 0.36 ± 1.31) (Fig. 4a).

**Figure 4:**
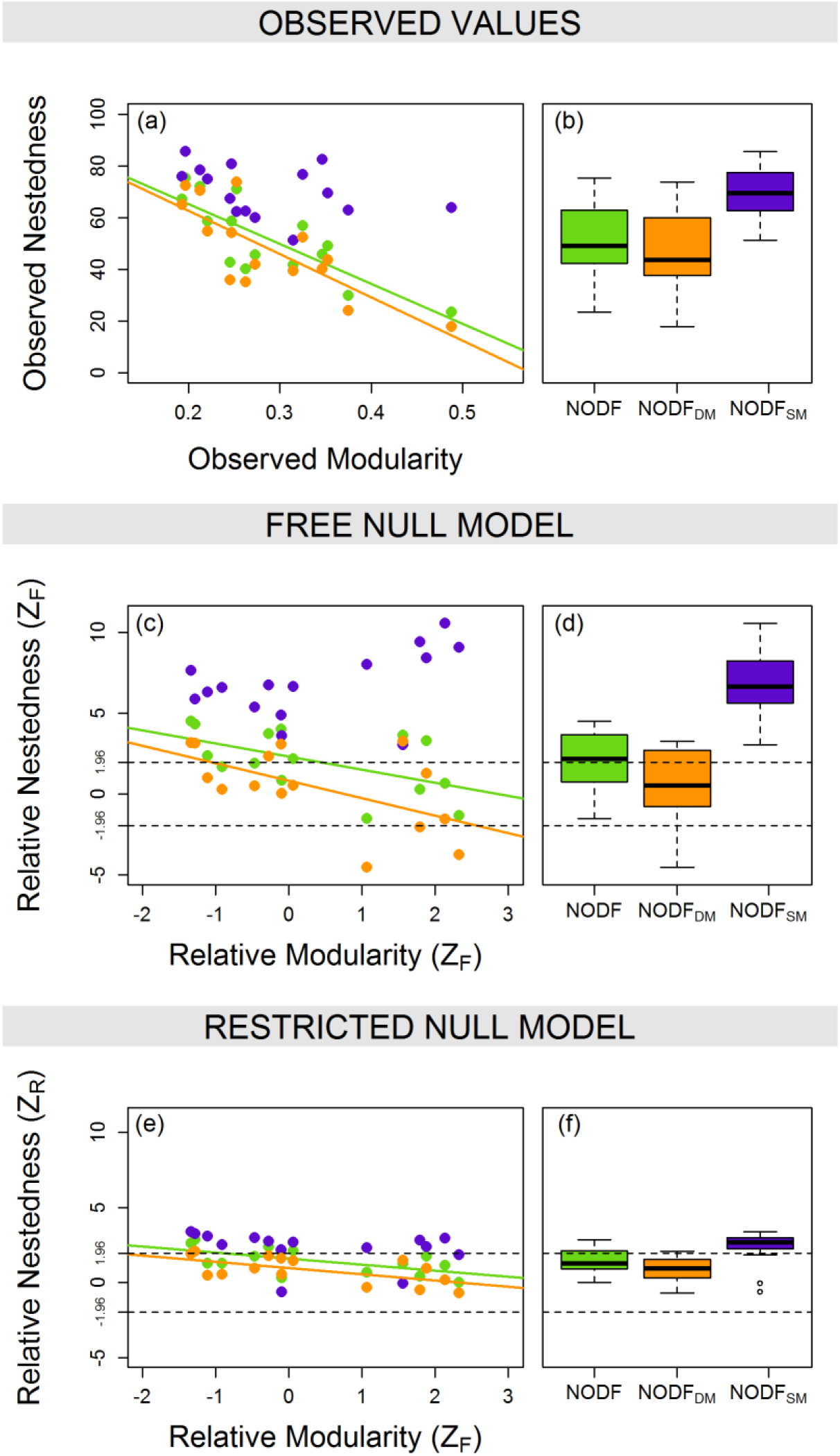
Relationship between observed and relative (Z_F_ and Z_R_) scores of nestedness components (NODF, NODF_SM_ and NODF_DM_) and modularity in local networks. Z_F_ and Z_R_ represent the relative score of a metric (nestedness or modularity) standardized by the score expected in the absence (the free null model) or in the presence (restricted null model) of the modular structure, respectively. NODF (green): overall nestedness. NODF_SM_ (purple): nestedness between pairs of species of the same module. NODF_DM_ (orange): nestedness between pairs of species of different modules. Left panels (a, c and e): relationship between observed and relative nestedness and modularity scores. Right panels (b, d and f): box plots of observed and relative nestedness scores. Notice that relative modularity is always standardized by modularity expected by the free null model, both in c and e. Lines show significant relationships (p<0.05).

However, some relationships between these two topologies were evident. First, both observed and relative overall nestedness (NODF) (Fig. 4a, c and e; and Appendix S2: Table S2) decreased as modularity increased. Second, the same pattern was true for nestedness between pairs of species of different modules (NODF_DM_) (Fig. 4a,c and e; and Appendix S2: Table S2). Third, although nestedness between pairs of species of the same module (NODF_SM_) was not significantly related to modularity in any case (Appendix S2: Table S2), the Z_F_NODF_SM_ showed a trend to increase with Z_F_Q (Fig. 4c).

In addition, as for the global networks, observed and relative values of NODF_SM_ were higher than those of NODF_DM_ (Fig. 4b,d and f), and the difference between them increased with modularity. Finally, NODF_SM_ values were higher than expected by both null models, while NODF_DM_ values were smaller than expected by the free null model but equal to that expected by the restricted null model (Fig. 4b,d and f).

### Specialization versus performance

Random factors explained a significant portion of the variance in flea abundance (Appendix S2: Table S3). In addition, as expected, only the within module degree (Z), participation coefficient (P), and the interaction between P and Z_F_NODF_DM_ significantly affects flea abundances. Neither Z_F_NODF_DM_ nor its interaction with Z was retained in the minimum selected model (Appendix S2: Table S4 and Table S5).

On the one hand, flea abundances were always positively correlated with within-module degree (Z) (Fig. 5). On the other hand, as expected, the relationship between flea abundance and participation coefficient (P) changed from positive to negative as the NODF_DM_ becomes smaller than expected by the free null model, crossing zero at Z_F_NODF_DM_ ≈ −2 (Fig. 5). In addition, the predicted positive effect of participation coefficient (P) on flea performance was higher than that of within-module degree (Z) when NODF_DM_ becomes equal to or higher than expected by the free null model.

**Figure 5:**
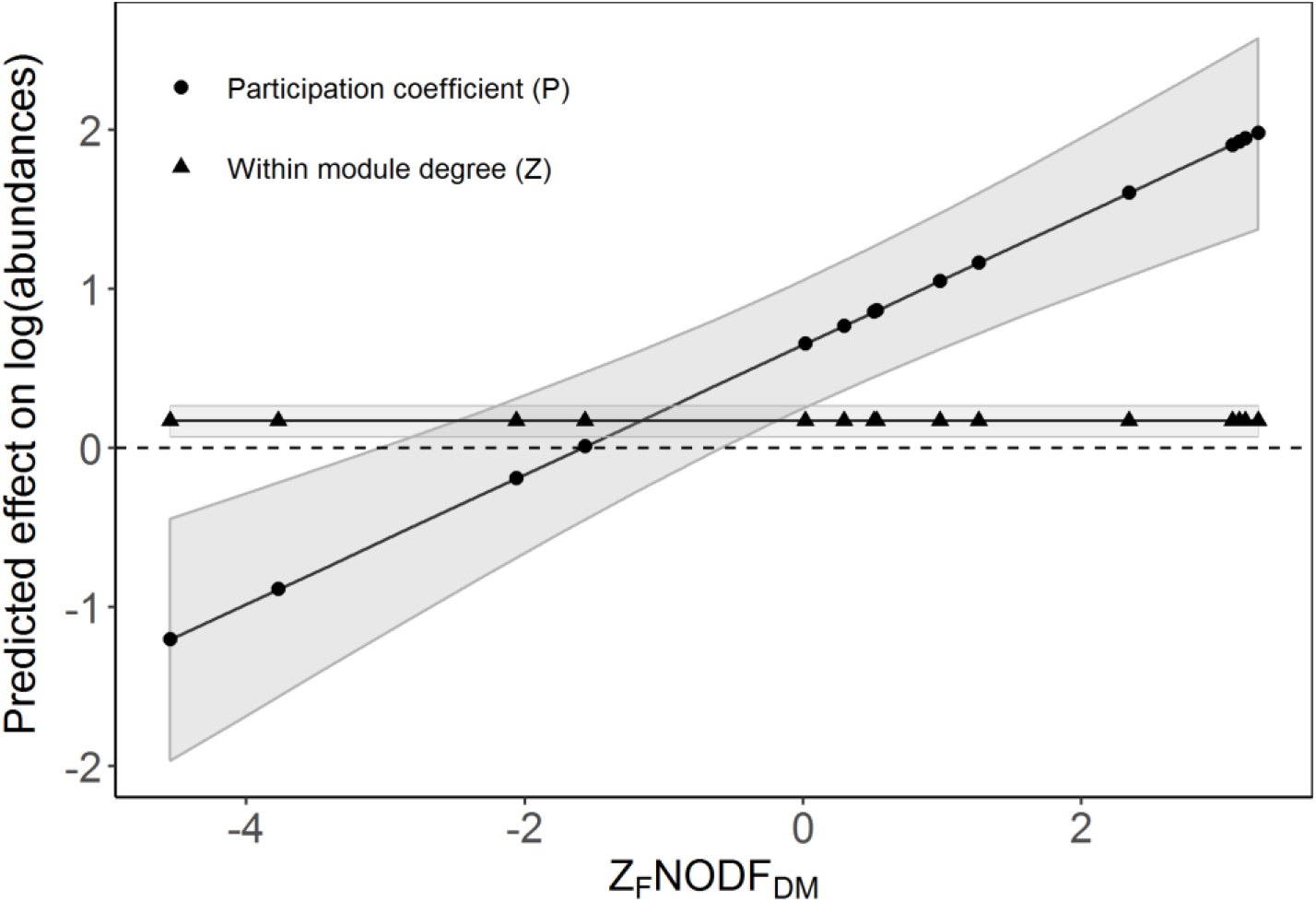
The predicted effects of within-module degree (Z) and participation coefficient (P) on flea performances (abundance) for different values of Z_F_NODF_DM_ (relative nestedness between pairs of species at different modules in a given local network when compared to that expected by the restricted null model). Lines show the predicted effects of P and Z on abundance (and its 95% confidence intervals), while dots indicate the fifteen local networks. As expected by the IHS, the effect of P changes from negative to positive (becoming higher than the positive effect of Z) as nestedness between species in different modules increases. The effect of Z on abundance in a given local network is independent of Z_F_NODF_DM_.

Finally, although the complete models explained a large amount of data variance, the fixed factors were responsible for only a very small fraction of the explanation (R^2^_(*m*)_ = 0.086, R^2^_(*c*)_ = 0.53).

## DISCUSSION

In the present study, we provide strong support for the integrative hypothesis of specialization (Pinheiro *et al*. 2016) using a continent-wide host-parasite network. We confirmed both (i) the emergence of a compound topology in the local and global networks (Fig. 2, 3 and 4); and (ii) the scale-dependence of the relationship between specialization and performance (Fig. 5). Our results unite two long-standing debates in the ecological literature within the same theoretical framework and provide clues to their solution.

### Nestedness versus modularity

What is the predominant topology in ecological networks: nestedness or modularity? In the past decades, conflicting results have been reported, leaving us with three possible scenarios. First, different systems, taxa, and interaction types lead to networks with different topologies. For instance, it has already been suggested that antagonistic networks tend to be modular, while mutualistic networks tend to be nested (Thebault & Fontaine 2010). Second, both topologies coexist as two sides of the same coin (Fortuna *et al*. 2010), with the nested structure superimposed over the modular structure. Third, alternative topologies should become manifest at different network scales, resulting in a compound topology (Lewinsohn *et al*. 2006). Our results provide strong support for the third scenario.

At first glance – considering only that the global and some local flea-mammal networks presented scores of nestedness and modularity higher than or equal to those expected by their species degrees (free null model) (Fig. 2a and Fig. 4c–d) – one could conclude that those two topologies coexist in the flea-mammal networks as two emergent properties of the same underlying phenomenon. However, the observed and relative values of modularity and overall nestedness were negatively correlated in the local networks (Fig. 4a,c and e). And while nestedness between pairs of species of the same module was higher than expected by the species degrees (NODF_SM_), the opposite was true for nestedness between pairs of species of different modules (NODF_DM_) (Fig. 4d). In addition, the difference between these two sets of nestedness increased with modularity (Fig. 4a,c and e). Together, those results strongly suggest that modularity constrains nestedness at large topological scales of this host-parasite system.

However, as pointed out in the Methods section, to say that the modules constrain nestedness between species of different modules and increase nestedness between species of the same module is a truism, a logical consequence of the definition of modules itself. In ecological terms, it is like saying that species preferences (specialization) constrain resource breadth processes (Brown 1984), which is trivial. A much more interesting issue would be to evaluate if interactions are more nested than expected *given* the species preferences, which are reflected in the network modular structure. This raises two questions.

First, is nestedness between pairs of species of the same module higher than expected *given* that they have similar dietary preferences, *i.e*, that they belong to the same module? By comparing the observed NODF_SM_ to that expected by the species degrees in the presence of a modular structure (the restricted null model), we showed that this is true for the flea-mammal network, which is formed by internally nested modules (Fig. 3).

Second, is nestedness between pairs of species of different modules higher than expected given that they have dissimilar dietary preferences, *i.e*, that they belong to different modules? That is, when a consumer *c_i_* of module *A* pervades the modular structure and consumes resources from module *B*, will *c_i_* consume the most consumed resources of module *B*? By comparing NODF_DM_ to that expected by the restricted null model, we show that this is also the case in the studied host-parasite system, since both in the global (Fig. 2b) and the local networks (Fig. 4f) the interactions between species of different modules were equally nested as expected by the restricted null model.

This scenario (NODF_SM_ and NODF_DM_ equal to or higher than expected given the modular structure) suggests that, once the constraints imposed by modules are overcome, the same processes that structure interactions within modules also structure the few interactions outside them. This does not necessarily need to be true, since competitive exclusion would predominate over resource availabilities outside the modules. For example, if parasites have poor performances in hosts that do not belong to their modules, diffused competition would prohibit parasites of module *B* to successfully establish themselves in the most exploited hosts of module *A*. If this happened, parasites would become supertramps (Diamond 1975) when exploiting hosts of other modules, and an antinested pattern of interactions would emerge between pairs of species of different modules (Poulin & Guégan 2000). Although it is not true in the flea-mammal network, it is an interesting question to be addressed in other systems.

Understanding the processes governing the spillover of species interactions between modules has important practical implications. It may help, for example, to predict which species are the most likely to invade new habitats or which parasite species are most likely to emerge in new hosts (Pimm 1991).

### The hypotheses of trade-offs and resource breadth

What is the expected relationship between the resource range of a species and its average performance at exploiting these resources (Futuyma & Moreno 1988)? As in the case of network topology, conflicting results have been reported, and two scenarios are possible. First, the relationship between resource breadth and average performance varies among systems, taxa, places, and interaction types. Second, this relationship should change at different community scales, from negative at larger scales to positive at smaller scales, as predicted by the IHS (Pinheiro *et al*. 2016).

Our results provide strong support for the second scenario (Fig 5). In local networks where modules represent a significant restriction to interactions, the relationship between flea abundance and generalism changed from positive within to negative among modules, as predicted by the IHS. In addition, the effect of *P* on abundance became more negative as the modular structure imposed more constraints on the interactions. Therefore, if the community (or, at least, the part we sampled) is composed of more than one module, a multi-scale relationship between specialization and performance should emerge. Otherwise, if the community is composed of very similar resources (*i.e*., just one internally nested module), we expect a simple positive relationship between generalism and performance. A further step would be to test if the contradictory results reported by previous studies which addressed the relationship between performance and generalism would also be explained by differences in the scale of each community studied. While some of them focused on different populations of the same resource species (e.g, Szollõsi *et al*. 2011) others sampled entire resource communities (Poulin 1998; Hellgren *et al*. 2009).

Fort *et al*. (2016) showed that higher abundance implies greater generalism in ecological networks, and not the contrary (but see Dorman *et al*. (2017)). That is, abundant species (the most available ones) are generalist because they have higher probability of finding potential interaction partners. This makes sense in the context of the IHS: the species with higher performances in exploiting a given set of similar resources reach higher abundances and, then, interact freely with a large number of that resources in the absence of trade-offs. Alternatively, although the nestedness between pairs of species of different modules was equal to the expected one *given* the modular structure in most of the local networks, the negative relationship between *P* and abundance suggests that generalism between modules has a negative effect on abundance, due to trade-offs. Therefore, although a species that is a hub in its own module would eventually spill over on the most connected species of other modules, if this species evolves adaptations to interact with a broad spectrum of dissimilar species, which makes it a hub in the entire network, this would have a negative impact on its performance.

### Concluding remarks

Some authors have pointed out that discontinuities are much more common in nature than previously thought. In addition, recent theoretical models (Holt 2006; Scheffer & van Nes 2006) suggest that, contrary to the principle of limiting similarity (Macarthur & Levins 1967), a balance between neutral and niche processes might generate self-organized clusters of similar species, which have been called emergent groups (Hérault 2007). The IHS and the compound topology are in complete agreement with this view. In fact, the idea that trade-offs generate modules, while a coupling between the availabilities of resources and consumers determines the dynamics inside each module, generating nestedness, suggests that there is a balance between niche processes at large scales and neutral processes at small scales.

Finally, we are not proposing that every ecological network should, necessarily, have a compound topology. Although it is likely that all nested matrices are modular at a larger scale, because any interactions should be constrained at some point, not all modules need to be internally nested. Several other patterns might be observed within a module, each resulting from different underlying mechanisms (Presley *et al*. 2010). The methods and the conceptual framework developed in our study provide tools to investigate these patterns.

### BOX 1 - The integrative hypothesis of specialization

Earlier coined the “integrative hypothesis of parasite specialization” (IHPS) (Pinheiro *et al*. 2016), the hypothesis was proposed in the context of host-parasite interactions as a solution to an old controversy in the parasitological literature: what is the expected relationship between generalism and performance of parasites (Poulin 2007)?

However, since its assumptions are broad enough to apply to other interactions systems, we describe it here as valid for any consumer-resource system and call it “the integrative hypothesis of specialization” (IHS). The IHS states that consumer-resource interactions are structured by a balance between resource breadth processes (Brown 1984) at small and trade-offs at large community scales.

On the one hand, at small community scales, resources should be very similar to one another and an adaptation (the arrow in Fig. 1a) that increases the intrinsic performance of a consumer in exploiting a specific resource type should also be an adaptation to all other resource types (the “+” in Fig 1a). Therefore, we do not expect to find preferences (e.g, phylogenetic, phenetic, or geographic signals) in consumer-resource interactions at small scales. Instead, a coupling between resource availabilities and intrinsic consumer performances should produce a nested pattern of realized performances: the resources with highest availability should be more strongly exploited by all consumers, in proportion to the intrinsic performances of the consumers on those resources (Fig. 1b).

Consequently, by sampling small scales of a community, one should find both a nested pattern of interactions (Fig. 1c), and a positive relationship between generalism and performance (Fig. 1d). This last pattern (Fig. 1d) is similar to the positive occupancy-abundance relationships widely reported for biogeographic data (Gaston & Blackburn 2000), normally explained by Brown’s resource breadth hypothesis (Brown 1984). Since the mechanisms described above are very similar to that proposed by Brown, but adapted to a context of interactions, we summarize them as resource breadth processes.

On the other hand, at larger community scales, the IHS states that there are trade-offs in the capacity to exploit resources of different clusters. Specifically, adaptations to a specific resource type (the arrow in Fig. 1e) should also be adaptations to other similar resource types of the same cluster (the “+” in Fig. 1e), but maladaptations to dissimilar resources of other clusters (the “-” in Fig. 1e). Thus, the pattern of realized performances described above (Fig. 1 b) should be restricted to within each resource cluster (Fig. 1f). In this context, the different clusters of resources are the real units of specialization and the true generalist consumers are those that can exploit resources at several clusters (blue species in figure 1). Consequently, by sampling large scales of a community, we do not expect to find a completely nested network, but rather a modular network with internally nested modules (i.e., a *compound topology*) (Lewinsohn *et al*. 2006) (Fig. 1g). In addition, although we expect a positive relationship between performance and generalism within each cluster of similar resources, we also expect trade-offs to result in a negative relationship between performance and capacity to exploit resources of different clusters (modules in the network) (Fig. 1h).

Krasnov *et al*. (2004) have already suggested that the relationship between performance and generalism should be negative in communities composed of dissimilar resources, but positive in communities composed of similar resources. Indeed, Krasnov et al’s suggestion and the IHS have the same underlying rationale: the probability of an adaptation to a given resource of being also an adaptation to other resources (i.e, the resource breadth hypothesis) is higher the more similar the resources are. The IHS is just a more inclusive hypothesis that also predicts the relationship between specialization and performance in communities in which resource dissimilarity is not gradually structured, that is, in communities with clusters of similar resources separated from one another by gaps of dissimilarity (Allen 2006).

## SUPPLEMENTARY FILES

### Appendix S1

Details of both free and restricted null models computations.

### Appendix S2

Supplementary Tables S1-S5

## ACKNOWLEDGEMENTS

We thank our institutions and many colleagues, who helped us in different ways during this project. Elisabeth Kalko, Judith Bronstein, Pedro Jordano, Nico Blüthgen, Carsten Dormann, Thomas Lewinsohn, Leonardo Rê-Jorge, Cang Hui, Adriano Paglia, Fernando Silveira and Erika Braga helped us with exciting discussions about ecological interactions, complex networks, and community assembly rules. We thank specially Prof. Thomas Lewinsohn for so many uplifting conversations about network topologies, which contributed substantially to conceptual and methodological issues of this work. The Graduate School in Ecology of the Federal University of Minais Gerais (ECMVS) provided us with scholarships granted to G.M.F. and R.B.P.P., and with infrastructure. M.A.R.M. was funded by the Alexander von Humboldt Foundation (AvH: 3.4-8151/15037), Minas Gerais Research Foundation (FAPEMIG: PPM-00324-15), Research Dean of the Federal University of Minas Gerais (UFMG-PRPq: 02/2014), Brazilian Council for Scientific and Technological Development (CNPq: 472372/2013-0), Brazilian Coordination for the Improvement of Higher Education Personnel (CAPES, grad student scholarships), and Research Program of the Biodiversity of the 14 Atlantic Forest (PPBio-MA/CNPq: 457458/2012-7).

